# A positive feedback loop drives centrosome maturation in flies

**DOI:** 10.1101/380907

**Authors:** Ines Alvarez-Rodrigo, Paul T. Conduit, Janina Baumbach, Zsofia A. Novak, Mustafa G. Aydogan, Alan Wainman, Jordan W. Raff

## Abstract

Centrosomes are formed when mother centrioles recruit pericentriolar material (PCM) around themselves. The PCM expands dramatically as cells prepare to enter mitosis (a process termed centrosome maturation), but it is unclear how this expansion is achieved. In flies, Spd-2 and Cnn form an extensive scaffold around the mother centriole that recruits other components of the mitotic PCM, and the Polo-dependent phosphorylation of Cnn at the centrosome is crucial for scaffold assembly. Here we show that, like Cnn, Spd-2 is specifically phosphorylated at centrosomes. This phosphorylation appears to create multiple phosphorylated S-S/T(p) motifs that allow Spd-2 to recruit Polo to the expanding scaffold. If Spd-2 cannot recruit Polo to the expanding scaffold, the scaffold is initially assembled around the mother centriole, but it cannot expand outwards, and centrosome maturation fails. We conclude that Spd-2, Polo and Cnn cooperate to form a positive feedback loop that drives the dramatic expansion of the mitotic centrosome in flies.

## Introduction

Centrosomes play an important part in many aspects of cell organisation, and they form when a mother centriole recruits pericentriolar material (PCM) around itself (Conduit et al., 2015). The PCM contains several hundred proteins (Alves-Cruzeiro et al., 2013), allowing the centrosome to function as a major microtubule (MT) organising centre, and also as an important coordination centre and signalling hub (Chavali et al., 2014; Vertii et al., 2016). Centrosome dysfunction has been linked to several human diseases and developmental disorders, including cancer, microcephaly and dwarfism (Bettencourt-Dias et al., 2011; Nigg and Raff, 2009).

During interphase, the mother centriole recruits a small amount of PCM that is highly organised (Lawo et al., 2012; Mennella et al., 2012; Sonnen et al., 2012; Fu and Glover, 2012). As cells prepare to enter mitosis, the PCM expands dramatically around the mother centriole in a process termed centrosome maturation (Palazzo et al., 2000). Electron microscopy (EM) studies suggest that centrioles organise an extensive “scaffold” structure during mitosis that surrounds the mother centriole and recruits other PCM components such as the γ-tubulin ring complex (γ-TuRC) (Moritz et al., 1995; 1998; Schnackenberg et al., 1998).

In the fruit fly *Drosophila melanogaster* and the nematode *Caenorhabditis elegans*, a relatively simple pathway seems to govern the assembly of this mitotic pericentriolar-scaffold. The conserved centriole/centrosome protein Spd-2/SPD-2 (fly/worm nomenclature) cooperates with a large, predominantly predicted-coiled-coil, protein (Cnn in flies, SPD-5 in worms) to form a scaffold whose assembly is stimulated by the phosphorylation of Cnn/SPD-5 by the mitotic protein kinase Polo/PLK-1 (Conduit et al., 2014a; 2014b; Feng et al., 2017; Woodruff et al., 2017; 2015; Wueseke et al., 2016). Mitotic centrosome maturation is abolished in the absence of this pathway, and some aspects of Cnn and SPD-5 scaffold assembly have recently been reconstituted *in vitro* (Feng et al., 2017; Woodruff et al., 2015; 2017). Vertebrate homologues of Spd-2 (Cep192) (Gomez-Ferreria et al., 2007; Zhu et al., 2008), Cnn (Cdk5Rap2/Cep215) (Barr et al., 2010; Choi et al., 2010; Fong et al., 2008; Kim and Rhee, 2014; Lizarraga et al., 2010) and Polo (Plk1) (Haren et al., 2009; Lane and Nigg, 1996; Lee and Rhee, 2011) also have important roles in mitotic centrosome assembly, indicating that elements of this pathway are likely to be conserved in higher metazoans. In vertebrate cells another centriole and PCM protein, Pericentrin, also has an important role in mitotic centrosome assembly that is dependent upon its phosphorylation by Plk1 (Haren et al., 2009; Lee and Rhee, 2011). In flies, the Pericentrin-like-protein (PLP) has a clear, but relatively minor, role in mitotic PCM assembly when compared to Spd-2 and Cnn (Lerit et al., 2015; Martinez-Campos et al., 2004; Richens et al., 2015).

Although most of the main players in mitotic centrosome-scaffold assembly appear to have been identified, several fundamental aspects of the assembly process remain unexplained. Cells entering mitosis, for example, contain two mother centrioles that assemble two mitotic centrosomes of equal size. It is unclear how this is achieved, as even a slight difference in the initial size of the two growing centrosomes would be expected to lead to asymmetric centrosome growth—as the larger centrosome would more efficiently compete for scaffolding subunits (Conduit et al., 2015; Zwicker et al., 2014). Fly centrioles appear to overcome this problem by constructing the mitotic centrosome-scaffold from the “inside-out”: in fly embryos, Spd-2 and Cnn are only incorporated into the scaffold close to the mother centriole, and they then flux outwards to form an expanded scaffold around the mother centriole (Conduit et al., 2014b; 2010). In this way, the pericentriolar scaffolds ultimately attain the same steady-state size—where incorporation around the mother centriole is balanced by loss of the scaffold at the centrosome periphery—irrespective of any initial size difference prior to mitosis (Conduit et al., 2015).

A potential problem with this “inside-out” mode of assembly is that the rate of centrosome growth is limited by the very small size of the centriole. Mathematical modelling indicates that the incorporation of a crucial PCM scaffolding component only around the mother centriole cannot easily account for the high rates of mitotic centrosome growth observed experimentally (Zwicker et al., 2014). To overcome this problem, it has been proposed that centrosome growth is “autocatalytic”, with the centriole initially recruiting a key organising component that can subsequently promote its own recruitment (Woodruff et al., 2014; Zwicker et al., 2014). Interestingly, it has been proposed that Spd-2 and Cnn could form a positive feedback loop that might serve such an autocatalytic function: Spd-2 helps recruit Cnn into the scaffold, while Cnn helps to maintain Spd-2 within the scaffold—so allowing higher levels of Spd-2 to accumulate around the mother centriole, which in turn drives higher rates of Cnn incorporation (Conduit et al., 2014b).

In worms (Decker et al., 2011) and vertebrates (Joukov et al., 2014; 2010; Meng et al., 2015) SPD-2/Cep192 can help recruit PLK1/Plk1 to centrosomes and, in vertebrates, Cep192 can also activate Plk1, in part through recruiting and activating Aurora A, another mitotic protein kinase implicated in centrosome maturation. We suspected, therefore, that in flies Spd-2 might recruit Polo into the centrosome-scaffold to phosphorylate Cnn and so help to generate a positive feedback loop that drives the expansion of the mitotic PCM. In flies, however, no interaction between Polo and Spd-2 has been reported yet. Indeed, an extensive Y2H screen for interactions between key centriole and centrosome proteins identified interactions between Spd-2 and the mitotic kinases Aurora A and Nek2, and between Polo and the centriole proteins Sas-4, Ana1 and Ana2, but not between Polo and Spd-2 (Galletta et al., 2016). A possible explanation for this result is that Polo/Plk1 is usually recruited to its interacting partners through its Polo-Box-Domain (PBD), which binds to phosphorylated S-S/T(p) motifs (Elia et al., 2003). Perhaps any Polo binding sites in fly Spd-2 were simply not phosphorylated in the Y2H experiments. In support of this possibility, a single phosphorylated S-S/T(p) motif in SPD-2 and Cep192 is required for these proteins to efficiently recruit PLK1/Plk1 to centrosomes in worms (Decker et al., 2011) and frogs (Joukov et al., 2010), while two such S-S/T(p) motifs (one of which is conserved with the single motif identified in frogs) have been identified in human Cep192 (Meng et al., 2015).

Here we examine the potential role of Spd-2 in recruiting Polo to centrosomes in *Drosophila* embryos. We show that, like Cnn, Spd-2 is largely unphosphorylated in the cytosol, but is highly phosphorylated at centrosomes, where Spd-2 and Polo extensively co-localise within the pericentriolar scaffold. We generate forms of Spd-2 that cannot bind the PBD of Polo by mutating S-S/T motifs in Spd-2 to T-S/T. These mutant Spd-2 proteins are still recruited to mother centrioles, where they can initiate the assembly of a centrosome scaffold with Polo and Cnn. Strikingly, however, the Spd-2/Polo/Cnn scaffold can no longer expand outwards around the mother centriole, and centrosome maturation fails. We conclude that Spd-2, Polo and Cnn form a positive feedback loop that drives the rapid expansion of the mitotic centrosome in flies.

## Results

### Spd-2 is phosphorylated specifically at centrosomes

We showed previously that Cnn is specifically phosphorylated at centrosomes (Conduit et al., 2014a), and we wondered if this was also the case for Spd-2. We partially purified centrosomes from embryo extracts by sucrose step-gradient centrifugation and compared the electrophoretic mobility of Spd-2 on western blots of the gradient fractions (Figure 1A). As was the case for Cnn, we observed a prominent slower migrating form of Spd-2 in the heavier centrosomal fractions that was largely absent in the lighter cytosolic fractions. However, unlike Cnn, a faster migrating form of Spd-2 was also present in the centrosomal fractions. Treatment of the centrosomal fractions with phosphatase revealed that the reduced mobility of Spd-2 in the centrosomal fractions could be attributed to phosphorylation (Figure 1B). Thus, like Cnn, Spd-2 is specifically phosphorylated at centrosomes, although not all of the Spd-2 at the centrosome appears to be phosphorylated.

**Figure 1.**
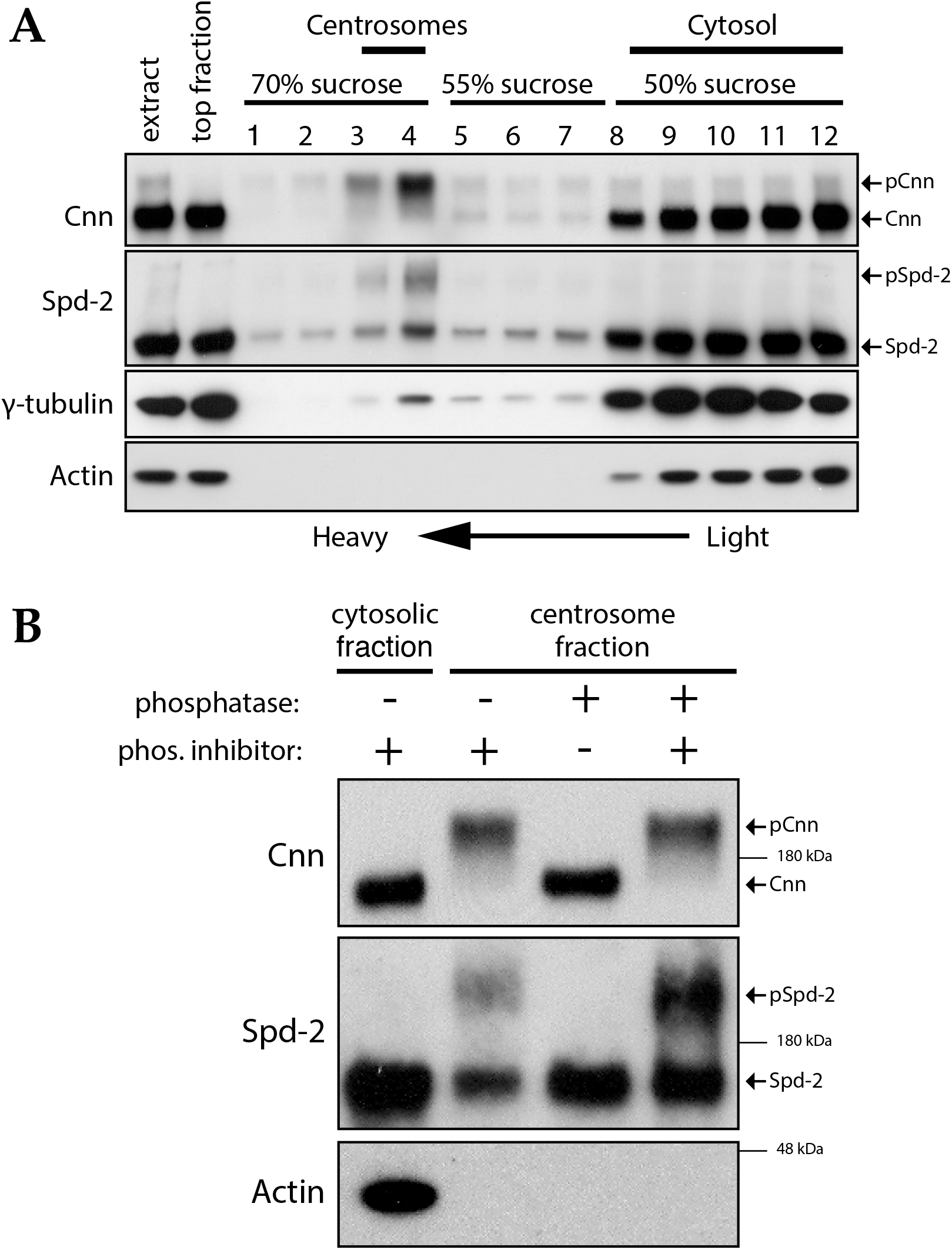
Spd-2 and Cnn are specifically phosphorylated at centrosomes. **(A)** Western blot of a sucrose step-gradient purification of centrosomes from embryo extracts probed with anti-Cnn, Spd-2, γ-tubulin and Actin antibodies, as indicated; sucrose steps are indicated above the blots. The gradient fractions are labelled 1 (heaviest) to 12 (lightest), and the cytosolic and centrosomal-peak fractions are indicated above the blot. Cnn, Spd-2 and γ-tubulin are enriched in the centrosomal fractions, where slower migrating species of Cnn and Spd-2, but not γ-tubulin, are detected. **(B)** A western blot of centrosomal fractions from the step gradient that were treated with phosphatase or phosphatase inhibitor (as indicated), and probed with the indicated antibodies; the cytosolic fraction is also shown. Note that the slower migrating species of Cnn and Spd-2 are absent after phosphatase treatment, but not if phosphatase inhibitor is included in the reaction. Note also that the Cnn and Actin blots shown in (B) are identical to the ones we presented in a previous publication (Conduit et al., 2014a), as they were performed contemporaneously with the Spd-2 blot shown here. Arrows indicate the identity of differently migrating species of the same protein. The blots shown are from one of two biological repeats that both gave similar results.

### Mutating multiple centrosomal phosphorylation sites in Spd-2 only mildly perturbs Spd-2 function *in vivo*

Using Mass Spectroscopy we identified seven Spd-2 peptides that were phosphorylated consistently and with high confidence in the centrosomal fractions, but not in the cytosolic fractions (Table S1). One of these peptides contained a phosphorylated S-S(p) motif, which could potentially help recruit Polo to centrosomes. We generated transgenic lines expressing WT Spd-2-GFP and a mutant form of Spd-2-GFP in which all 7 of the centrosomally phosphorylated residues—together with an additional 4 Ser/Thr resides that could potentially be phosphorylated in these peptides (Table S1)—were mutated to Ala (Spd-2-11A-GFP). Interestingly, although Spd-2-11A-GFP was expressed at substantially lower levels than WT Spd-2-GFP (Figure S1A), both Spd-2-GFP and Spd-2-11A-GFP rescued the female sterility phenotype of *Spd-2* mutant flies, and in *Spd-2* mutant embryos the mutant protein localised to centrosomes nearly as well as the WT protein (Figure S1B-D). Thus, although present at lower levels, Spd-2-11A-GFP is strongly recruited to centrosomes, and preventing the phosphorylation of at least 7 sites in Spd-2 that are specifically phosphorylated at centrosomes only mildly perturbs Spd-2 function *in vivo*.

### Generating a form of Spd-2 that should not be able to recruit Polo

The Spd-2 phosphorylation sites we identified here were also identified previously in several phospho-proteomic screens in *Drosophila melanogaster* (Bodenmiller et al., 2007; Habermann et al., 2012; Zhai et al., 2008). These screens identified 11 additional Spd-2 peptides that each contained at least one phosphorylated S-S/T(p) motif that could potentially recruit Polo to centrosomes (summarised in Table S2).

We wondered, therefore, whether multiple S-S/T(p) motifs in *Drosophila* Spd-2 might help recruit Polo to centrosomes. To rigorously test this possibility, we generated transgenic lines expressing a Spd-2-GFP fusion in which all 34 S-S/T motifs in *Drosophila melanogaster* Spd-2 were mutated to T-S/T (**Spd-2-ALL-GFP**) (*blue* and *red* lines, Figure 2A,B). The conservative substitution of Thr for Ser should not disturb the structure of the protein or block the ability of this motif to be phosphorylated, but should prevent this phosphorylated motif from binding the PBD (Elia et al., 2003). In addition, 16 of the 34 S-S/T motifs were highly conserved in *Drosophila* species (Figure 2A, *red* lines), so we also generated transgenic lines expressing a form of Spd-2-GFP in which only these 16 conserved motifs were mutated (**Spd-2-CONS-GFP**). Based on our observations reported below, we refer to these mutant proteins as being unable to recruit Polo.

**Figure 2.**
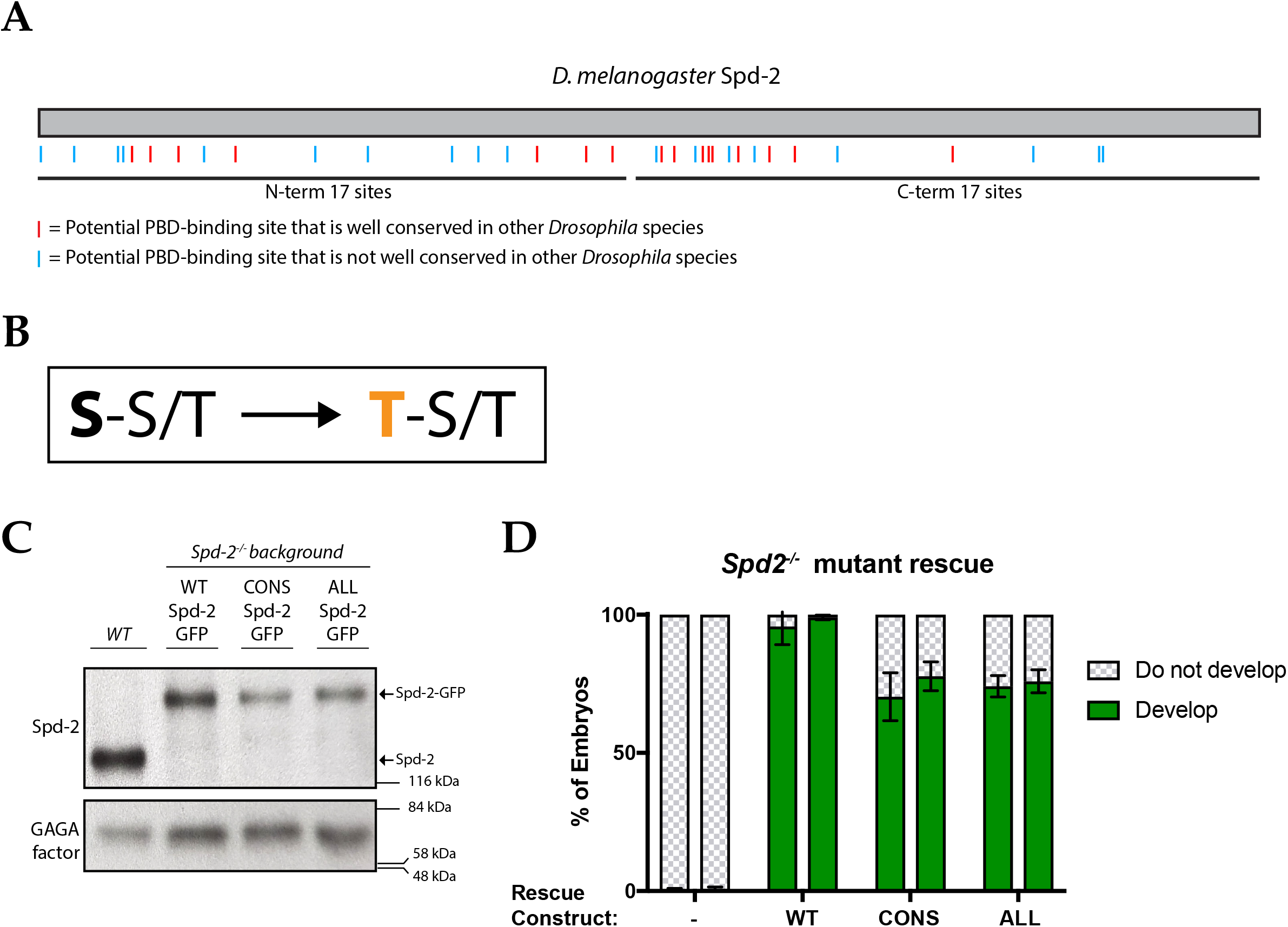
Generating mutant forms of Spd-2 that should not be able to bind to the PBD. **(A)** A schematic representation of the *Drosophila melanogaster Spd-2* amino acid sequence, indicating the position of S-S/T motifs (potential PBD-binding sites) that are either highly conserved (*red* lines), or not highly conserved (*blue* lines), in other *Drosophila* species. **(B)** Schematic of the S-S/T ➔ T-S/T mutations introduced in Spd-2-CONS-GFP and Spd-2-ALL-GFP. **(C)** A western blot of WT embryos, or *Spd-2* mutant embryos expressing either WT Spd-2-GFP, Spd-2-CONS-GFP or Spd-2-ALL-GFP transgenes, probed with anti-Spd-2 antibodies, or the GAGA transcription factor (Raff et al., 1994) as a loading control. The blot shown is from one of multiple technical replicates that gave similar results. **(D)** Bar charts quantify the percentage of *Spd-2* mutant embryos that had initiated development after expression of either the WT Spd-2-GFP, Spd-2-CONS-GFP or Spd-2-ALL-GFP transgenes (as indicated). The chart shows the data from two independent biological repeats; 3 lots of >50 embryos that were collected and processed independently were scored for each repeat; error bars represent the standard deviation (SD) for each repeat.

Western blotting revealed that the mutant Spd-2-GFP-fusions were expressed at slightly lower levels than WT Spd-2-GFP in embryos (Figure 2C)—although this difference was not as striking as that seen with Spd-2-11A-GFP (Figure S1A). Both mutant proteins significantly rescued the defect in pronuclear fusion in *Spd-2* mutant embryos (so allowing these embryos to start to develop)—although to a slightly lesser extent than the WT protein (Figure 2D). Unlike Spd-2-11A-GFP, however, *Spd-2* mutant embryos expressing either Spd-2-CONS-GFP (hereafter **Spd-2-CONS-GFP embryos**) or Spd-2-ALL-GFP (hereafter **Spd-2-ALL-GFP embryos**) almost never hatched as larvae (<1/500 embryos hatching; n>2000 embryos scored).

### Spd-2 that cannot recruit Polo is less concentrated at centrosomes, and these centrosomes organise less robust asters of MTs

To investigate why Spd-2-CONS-GFP and Spd-2-ALL-GFP embryos almost never hatched as larvae, we expressed a Jupiter-mCherry transgene in these embryos to follow the behaviour of MTs. WT Spd-2-GFP localised robustly to centrosomes at all stages of the embryonic cell cycle, and the centrosomes organised robust astral and spindle MT arrays (Figure 3A, left panels). The mutant proteins were less concentrated at centrosomes (Figure 3A,B) and the astral MT arrays organised by the centrosomes were less robust (Figure 3A,C). Embryos expressing the mutant proteins exhibited progressively more severe mitotic defects as they developed, and these defects were qualitatively, but reproducibly, more pronounced in Spd-2-ALL-GFP embryos (Figure 3A, *white* arrowheads). We conclude that the mutant Spd-2-GFP fusions can support the assembly of a centrosome that usually allows pronuclear fusion to occur (and so usually allows embryos to initiate development), but these centrosomes cannot support the very rapid syncytial mitotic divisions—and so most of these embryos die during early embryonic development.

**Figure 3.**
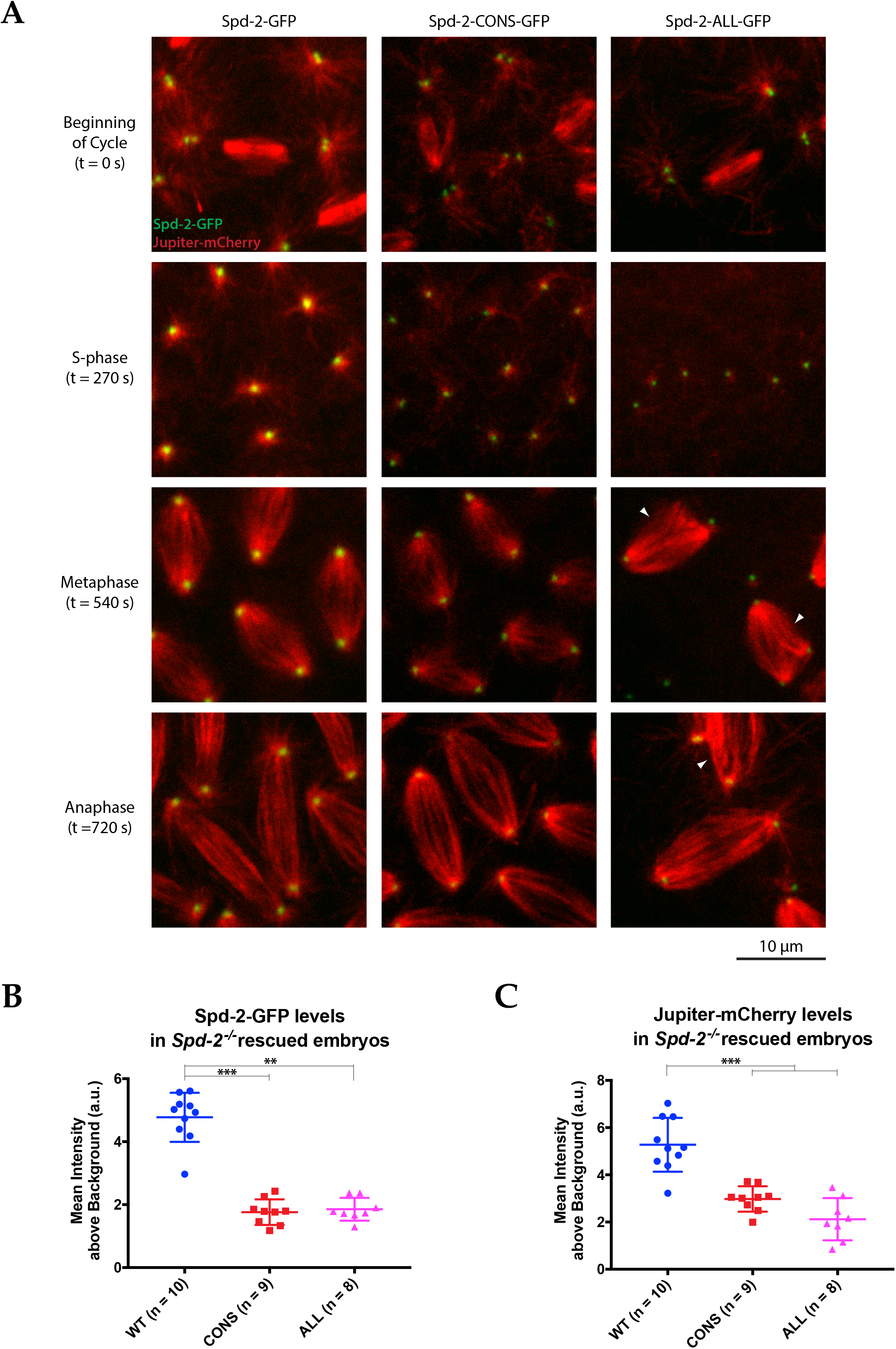
Centrosomes recruit less Spd-2 and organise fewer MTs if Spd-2 cannot bind Polo. **(A)** Micrographs show stills of living *Spd-2* mutant embryos expressing the MT-marker Jupiter-mCherry (*red*) and either WT Spd-2-GFP, Spd-2-CONS-GFP or Spd-2-ALL-GFP (*green*, as indicated). Time (in seconds) is indicated as the embryos progress from early S-phase (t=0s) to Anaphase (t=720s). Note that the Spd-2-ALL-GFP embryos are disorganised and exhibit relatively severe mitotic defects making them difficult to compare directly to the Spd-2-GFP embryos (*white* arrowheads). **(B,C)** Graphs quantify the centrosomal intensity of the various Spd-2-GFP transgenes (B) or the centrosomal MT intensity (C) during late S-phase (t=270s). Each dot represents the average intensity of the 5 brightest centrosomes in a single embryo; n=number of embryos analysed; error bars indicate the mean +/− SD of each population of embryos scored. The D’Agostino-Pearson omnibus normality test was used to check Gaussian distribution of the data. One-Way ANOVA with Tukey’s multiple comparisons test was used when data passed the normality test (Jupiter-mCherry datasets); Kruskal-Wallis test with Dunn’s multiple comparisons test was used otherwise (**, P<0.01; ***, P<0.001).

### Spd-2 that cannot recruit Polo is still recruited to centrioles, but it does not efficiently form a scaffold that spreads outwards

We next used live-cell 3D-structured illumination super-resolution microscopy (3D-SIM) to examine in more detail the centrosomal localisation of the Spd-2 mutant proteins in *Spd-2* mutant embryos. As shown previously (Conduit et al., 2014b), WT Spd-2-GFP localised to a toroid structure surrounding the mother centriole, and also to a fibrous scaffold-like structure that extended outwards around the mother centriole (Figure 4A, left panels). Strikingly, both mutant proteins still localised strongly to the mother centriole, but the amount of mutant proteins in the pericentriolar scaffold was greatly reduced—suggesting either that the mutant proteins were unable to efficiently incorporate into the scaffold, or that almost no scaffold was assembled in these embryos (Figure 4A).

**Figure 4.**
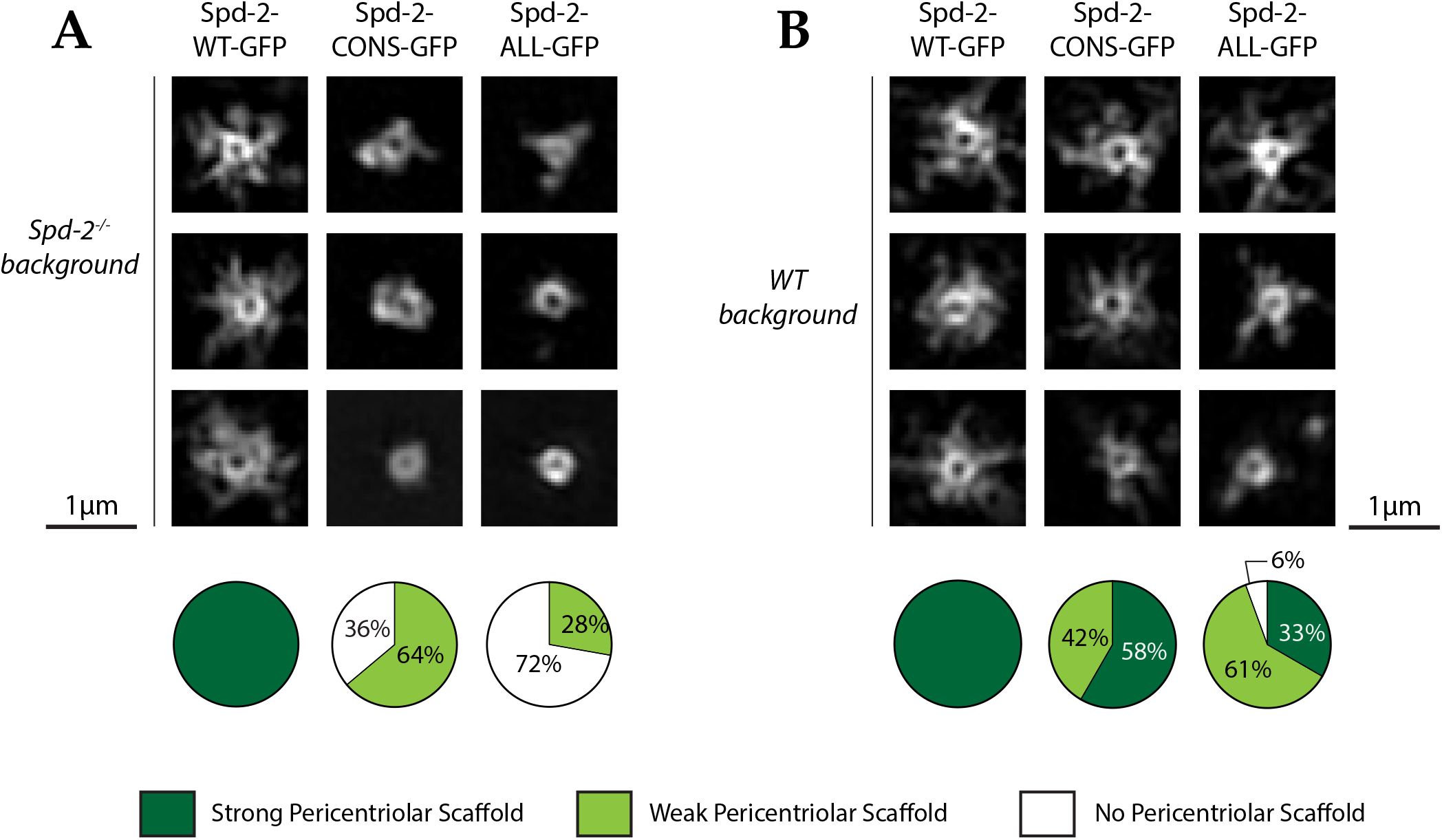
Spd-2 that cannot bind Polo does not efficiently form a pericentriolar scaffold. **(A,B)** Micrographs show 3D-SIM images of individual centrosomes from either *Spd-2* mutant embryos (A) or WT embryos (B) expressing WT Spd-2-GFP, Spd-2-CONS-GFP or Spd-2-ALL-GFP (as indicated). Pie charts underneath quantify the percentage of centrosomes that were scored qualitatively in blinded experiments as having a strong (*dark green*), weak (*light green*) or no (*white*) pericentriolar scaffold (n=36, for each genotype, respectively). Centrosome images were selected based on quality of the reconstruction as assessed by SIM-Check (Ball et al., 2015) and the presence of a visible, well-formed ring corresponding to the presence of protein at the mother centriole wall. Note that all centrosomes were imaged in mid-late S-phase when the centrosomal levels of Spd-2 are maximal (see below).

To test whether the mutant proteins could assemble into an existing pericentriolar scaffold, we expressed them in WT embryos that contain endogenous, unlabelled, Spd-2. Both mutant proteins could now incorporate into the scaffold, although their levels were reduced compared to WT Spd-2-GFP (Figure 4B; data not shown). The simplest interpretation of this data is that the WT and mutant proteins can co-assemble into a “mixed” pericentriolar scaffold, but the inability of the mutant proteins to recruit Polo reduces the assembly and/or maintenance of this mixed scaffold.

### The entire pericentriolar scaffold cannot efficiently expand outwards around the mother centriole if Spd-2 cannot recruit Polo

Polo and Cnn normally cooperate with Spd-2 to form the pericentriolar scaffold, so we tested whether Polo or Cnn could still efficiently form a scaffold even when the Spd-2-ALL and Spd-2-CONS proteins cannot. We examined the distribution of Polo-GFP in living *Spd-2* mutant embryos expressing either WT Spd-2-mCherry, or Spd-2-CONS-mCherry. Spd-2-ALL was excluded from this analysis because *Spd-2* mutant embryos co-expressing Polo-GFP and Spd-2-ALL-mCherry exhibited very severe developmental defects (Figure S2), and this was also true when Spd-2-ALL fusions were combined with several other GFP- or mCherry-fusions. WT Spd-2-mCherry and Polo-GFP extensively co-localised at the mother centriole and spread outwards together into the pericentriolar scaffold—supporting the idea that Spd-2 normally helps recruit Polo to this scaffold in flies (Figure 5A). In contrast, although the Spd-2-CONS-mCherry and Polo-GFP proteins still co-localised around the mother centriole, neither protein formed a robust pericentriolar scaffold (Figure 5B). As a result, the level of Polo-GFP at the centrosome was severely reduced. These results strongly indicate that phosphorylated S-S/T(p) motifs in Spd-2 are not required to recruit Polo to mother centrioles, but are required to allow Polo and Spd-2 to efficiently co-assemble into a scaffold around the mother centriole.

**Figure 5.**
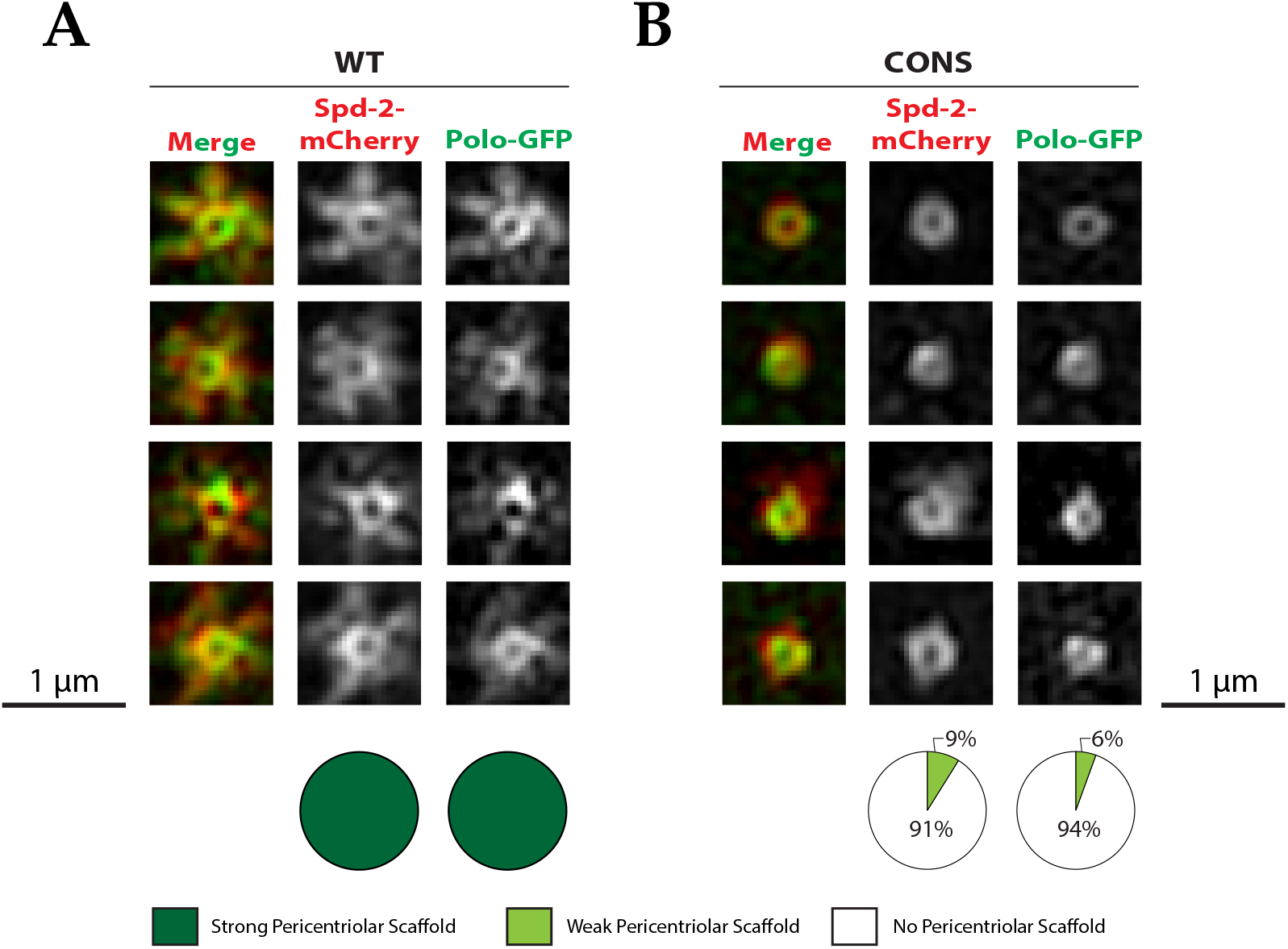
Polo is still recruited to the mother centriole, but it cannot assemble into a pericentriolar scaffold, if it cannot bind to Spd-2. Micrographs show 3D-SIM images of individual centrosomes from *Spd-2* mutant embryos expressing Polo-GFP (*green* in merged images) and either WT Spd-2-mCherry (A) or Spd-2-CONS-mCherry (B) (*red* in merged images). Pie charts quantify the percentage of centrosomes that were scored qualitatively in blinded experiments as having a strong (*dark green*), weak (*light green*) or no (*white*) scaffold for either Polo-GFP or Spd-2-mCherry scaffold (n=15 individual centrosomes, 2 images (channels) per centrosome, for each genotype, respectively). The full dataset was independently scored by 3 individuals, and the average score is shown. Note that the defect in pericentriolar scaffold assembly is slightly stronger in these *Spd-2* mutant embryos expressing Polo-GFP and Spd-2-CONS-mCherry than in the *Spd-2* mutant embryos expressing just Spd-2-CONS-GFP (Figure 4A). This appears to be due to a genetic interaction between Polo-GFP and Spd-2-CONS-mCherry, as mutant embryos expressing just Spd-2-CONS-mCherry had a similar phenotype to embryos expressing just Spd-2-CONS-GFP (data not shown), indicating that it is the co-expression of a GFP-tagged version of Polo together with an mCherry-tagged version of Spd-2-CONS that makes the defect in pericentriolar scaffold assembly more pronounced.

We next used 3D-SIM to examine the distribution of RFP-Cnn in living *Spd-2* mutant embryos expressing either WT Spd-2-GFP or Spd-2-CONS-GFP. In Spd-2-GFP embryos, RFP-Cnn spread outwards along the centrosomal MTs forming a robust pericentriolar scaffold that extended beyond the Spd-2-GFP scaffold (Figure 6A), as reported previously (Conduit et al., 2014b). In contrast, in Spd-2-CONS-GFP embryos only an occasional protrusion of RFP-Cnn and Spd-2-CONS-GFP extended outwards from the centriole (*white* arrowheads, Figure 6B), and both proteins generally remained tightly concentrated together around the mother centriole (Figure 6B). These relative distributions of RFP-Cnn were confirmed and quantified by “radial-profiling” using data obtained on a standard spinning disk confocal system (Conduit et al., 2014b) (Figure 6C). Strikingly, radial-profiling also revealed how the RFP-Cnn scaffold *(red* lines, Figure 6D) normally extends beyond the Spd-2-GFP scaffold (*green* lines in Figure 6D) in WT embryos, but these distributions essentially overlap in Spd-2-CONS-GFP embryos (Figure 6D). Thus, if Spd-2 cannot recruit Polo, the Cnn-scaffold cannot efficiently expand outwards beyond the Spd-2 scaffold.

**Figure 6.**
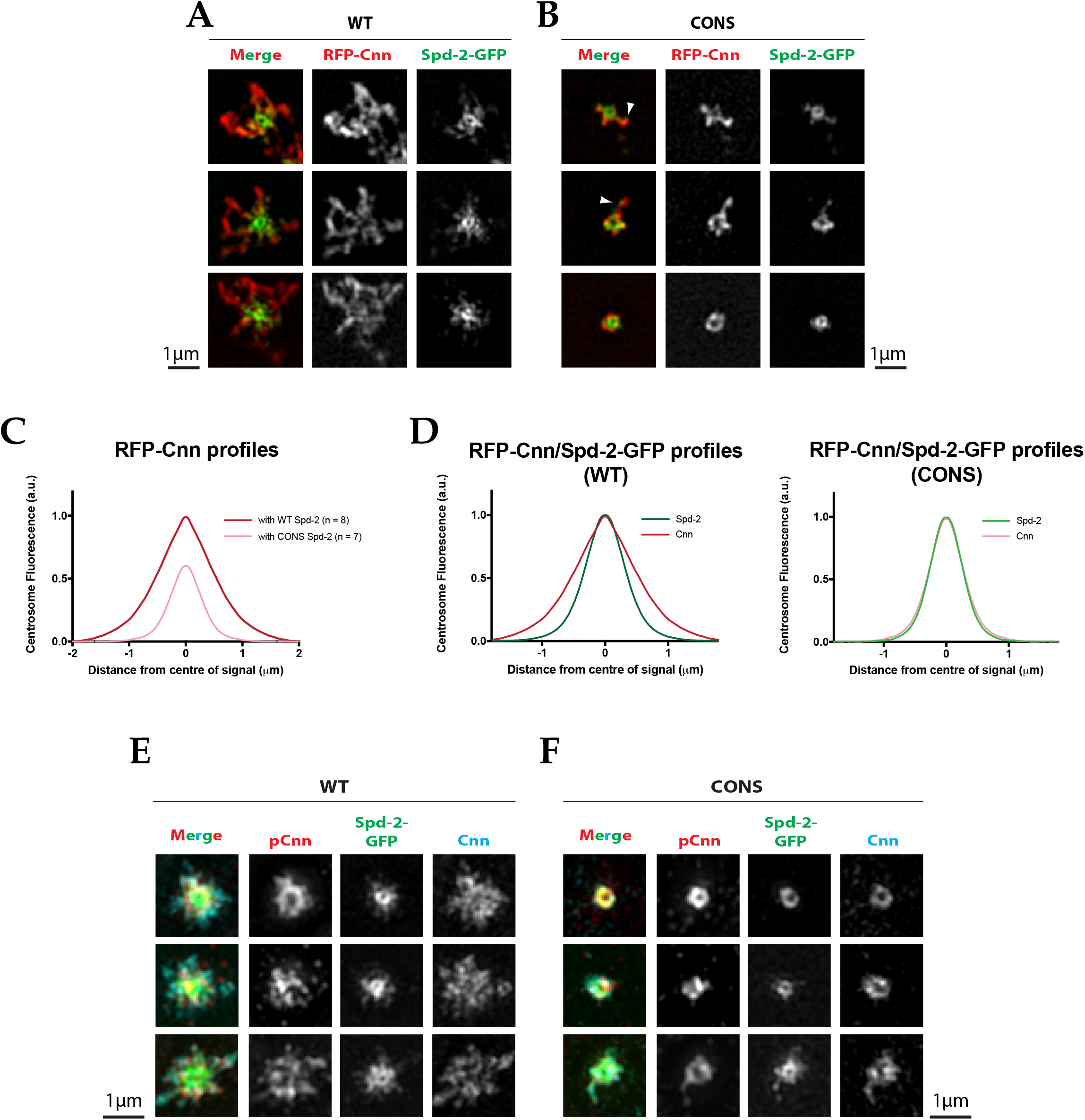
Cnn is still recruited to, and phosphorylated at, the mother centriole, but it does not efficiently assemble into a pericentriolar scaffold if Spd-2 cannot recruit Polo. **(A,B)** Micrographs show 3D-SIM images of individual centrosomes from *Spd-2* mutant embryos expressing RFP-Cnn (*red* in merged images) and either WT Spd-2-GFP (A) or Spd-2-CONS-GFP (B) (*green* in merged images). Arrowheads indicate examples of occasional protrusions of RFP-Cnn and Spd-2-CONS-GFP. **(C,D)** Graphs compare the radial distributions of RFP-Cnn around the mother centriole in WT Spd-2-GFP and Spd-2-CONS-GFP embryos (C), or the radial distribution of RFP-Cnn and Spd-2-GFP in either WT Spd-2-GFP or Spd-2-CONS-GFP embryos (D, as indicated). Data for these graphs was obtained from living embryos examined on a spinning disk confocal system; 7-8 embryos (as indicated), and 5 centrosomes in each embryo, were analysed. **(E,F)** Micrographs show 3D-SIM images of individual centrosomes from *Spd-2* mutant embryos expressing either WT Spd-2-GFP (E) or Spd-2-CONS-GFP (F) that were fixed and stained with antibodies against GFP, phospho-Cnn, or total-Cnn (*green, red* and *blue* in merged images, respectively).

Cnn is normally phosphorylated at centrosomes in a Polo-dependent manner, and this allows Cnn to assemble into a scaffold (Conduit et al., 2014a; Feng et al., 2017). To test whether the centrosomal Cnn could still be phosphorylated by Polo in Spd-2-CONS-GFP embryos, we stained Spd-2-GFP and Spd-2-CONS-GFP embryos with an antibody that specifically recognises a phospho-epitope in Cnn that is phosphorylated by Polo (Feng et al., 2017), as well as with antibodies that recognise the total Cnn protein (Figure 6E,F). Phosphorylated Cnn was still strongly detected around the mother centriole in Spd-2-CONS-GFP embryos, demonstrating that Cnn can still be phosphorylated at centrosomes even if Spd-2 cannot recruit Polo (Figure 6F). This phosphorylation is presumably dependent upon the Polo that is still recruited to the mother centriole in Spd-2-CONS-GFP embryos (Figure 5B). Thus, a functional Polo/Spd-2/Cnn “mini-scaffold” that contains phosphorylated Cnn and that can organise some MTs (Figure 3A) assembles around the mother centriole even if Spd-2 cannot itself recruit Polo; this scaffold, however, is unable to efficiently expand outwards around the mother centriole (Figure 5B; Figure 6B).

As the Polo/Spd-2/Cnn scaffold is essential for mitotic centrosome assembly in flies (Conduit et al., 2014b; Dobbelaere et al., 2008; Feng et al., 2017), the failure of these proteins to form an expanded pericentriolar scaffold in Spd-2-CONS embryos would be predicted to lead to a failure to recruit any other PCM proteins to an expanded mitotic PCM. This appeared to be the case, as the centrosomal recruitment of the PCM components Aurora A-GFP (Figure 7A-D) and γ-tubulin (Figure 7E-H) was greatly reduced in Spd-2-CONS-GFP embryos. Thus, the failure to form an expanded pericentriolar scaffold in these embryos appears to lead to a general failure in centrosome maturation.

**Figure 7.**
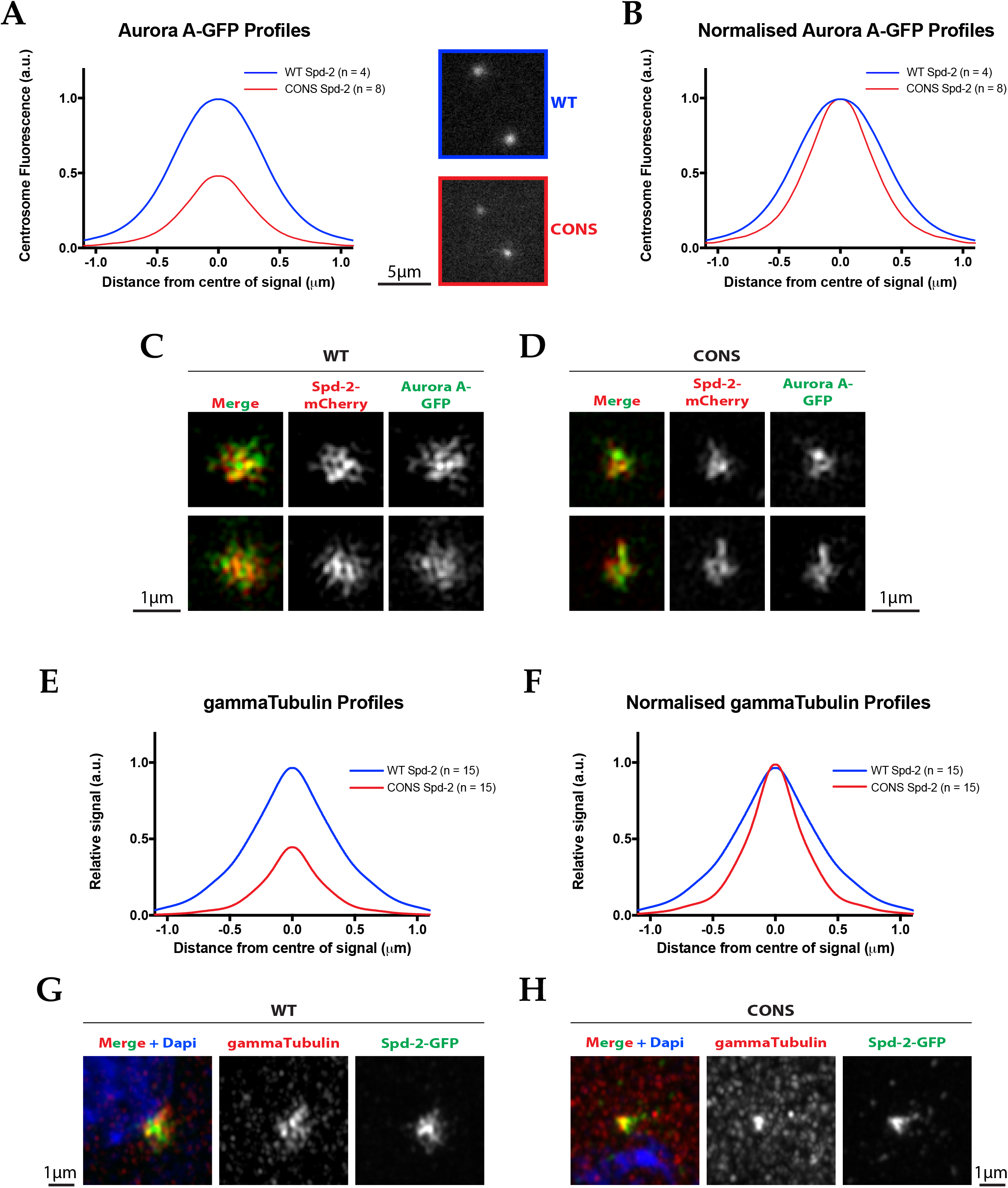
The mitotic PCM does not expand outwards around the mother centriole if Spd-2 cannot recruit Polo. **(A)** Graph compares the radial distribution of the PCM component Aurora-A-GFP around the mother centriole in living WT Spd-2-mCherry (*blue*) and Spd-2-CONS-mCherry embryos (*red*). Insets show examples of typical spinning disk-confocal images used for this analysis; 4-8 embryos (as indicated), and 5 centrosomes in each embryo, were analysed. **(B)** Graph shows the same data as shown in (A), but normalised so that the peak intensity of both genotypes = 1. This emphasises how even if the centrosomal Aurora A-GFP signal is normalised for fluorescence intensity, Aurora A-GFP spreads out around the mother centriole to a lesser extent in Spd-2-CONS-mCherry embryos than in WT Spd-2-mCherry embryos. **(C,D)** Micrographs show 3D-SIM images of individual centrosomes from *Spd-2* mutant embryos expressing Aurora A-GFP (*green* in merged images) and either WT Spd-2-mCherry (C) or Spd-2-CONS-mCherry (D) (*red* in merged images). **(E-H)** Same analysis as presented in panels (A-D), but analysing the distribution of the PCM component γ-tubulin in fixed embryos. For radial profiles, we analysed 1 pair of centrosomes per embryo, 5 embryos per technical replicate (embryos collected and processed independently), and 3 technical replicates in total per condition (total n=15).

### Centrosome maturation fails if Spd-2 cannot recruit Polo into the expanding mitotic PCM

To more directly examine the kinetics of centrosome maturation in Spd-2-CONS-GFP embryos we quantified the centrosomal levels of WT Spd-2-GFP and Spd-2-CONS-GFP through an entire embryonic cell-cycle. In *Drosophila*, the rapidly dividing syncytial embryos cycle between S- and M-phases with no intervening Gap periods—so as soon as the embryos enter S-phase the two centrosomes separate and start to mature in preparation for the next M-phase (Foe and Alberts, 1983). WT Spd-2-GFP started to accumulate at the maturing centrosomes in early S-phase, reached maximal levels just before nuclear envelope breakdown, and then started to decline as the embryos entered mitosis (*blue* line, Figure 8; Figure S3). In contrast, centrosomal levels of Spd-2-CONS-GFP remained at a constant low level throughout the cycle, indicating that centrosome maturation fails if Spd-2 cannot recruit Polo into the expanding pericentriolar scaffold. (*red* line, Figure 8; Figure S4).

**Figure 8.**
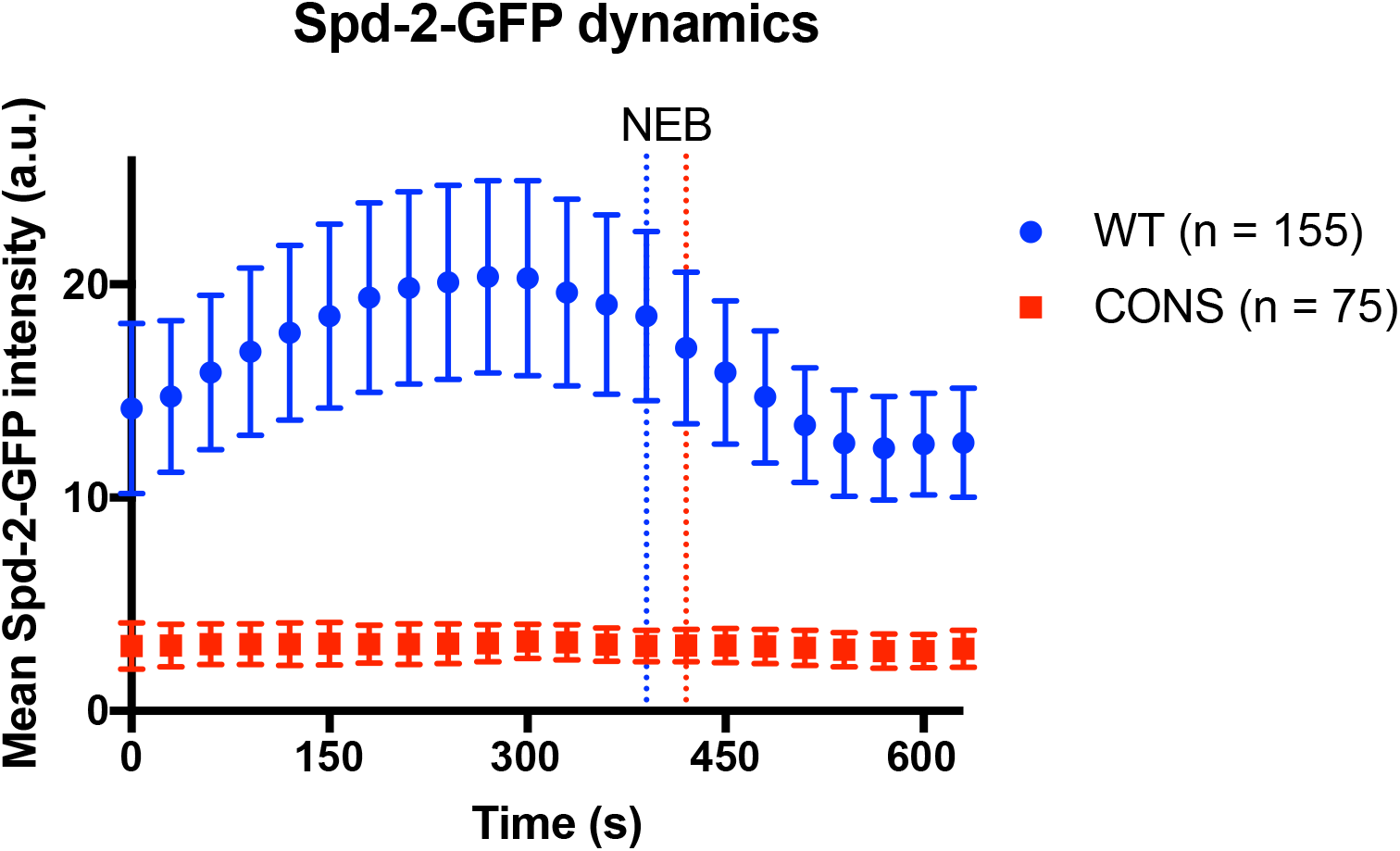
Centrosome maturation fails if Spd-2 cannot recruit Polo. Graph compares the mean centrosomal Spd-2-GFP intensity (in arbitrary units [a.u.]) through an entire embryonic cell cycle in either a WT Spd-2-GFP (*blue*) or Spd-2-CONS-GFP (*red*) embryo; n=number of centrosomes analysed per embryo. Bars indicate SD. Time in seconds is indicated, and the time when centrosomes first separate at the start of S-phase is set at t=0; the time of mitotic entry—scored as the time of nuclear envelope breakdown (NEB)—in each embryo is indicated by the dotted vertical lines. Because the length of S-phase varies slightly in each individual embryo—384s ± 46s or 369s ± 30s for WT Spd-2-GFP and Spd-2-GFP-CONS, respectively (average ± SD)—it is not possible to simply average the data at each time point from multiple embryos, so a representative example is shown here. Nevertheless, a similar pattern was observed in all 14 WT Spd-2-GFP (*blue*) or Spd-2-CONS-GFP (*red*) embryos that we monitored in this way (see Figure S3 and Figure S4).

### The ability to recruit Polo to the expanded mitotic PCM is not restricted to one specific site or region of Spd-2

Studies in worms and frog extracts have concluded that a single specific S-S/T(p) motif is required to allow SPD-2/Cep192 to efficiently recruit PLK-1/Plk1 to centrosomes, and in human cells a second specific S-S/T(p) motif also has an important role (Decker et al., 2011; Joukov et al., 2014; Meng et al., 2015). However, our data is consistent with the possibility that several, and perhaps up to 16, conserved S-S/T(p) motifs in *Drosophila* Spd-2 may help recruit Polo to centrosomes. Moreover, our observation that Spd-2-ALL-GFP embryos consistently exhibited more severe developmental abnormalities than the Spd-2-CONS-GFP embryos suggests that one or more non-conserved S-S/T(p) motifs may contribute to this process if the conserved sites are absent. To examine whether Polo recruitment by fly Spd-2 could be specifically linked to any of the previous S-S/T(p) motifs identified, we assayed the ability of several mutant Spd-2-mKate2 fusions to recruit Polo-GFP using a mRNA injection assay into WT embryos that we used previously to study this process (Novak et al., 2016) (Figure 9A).

**Figure 9.**
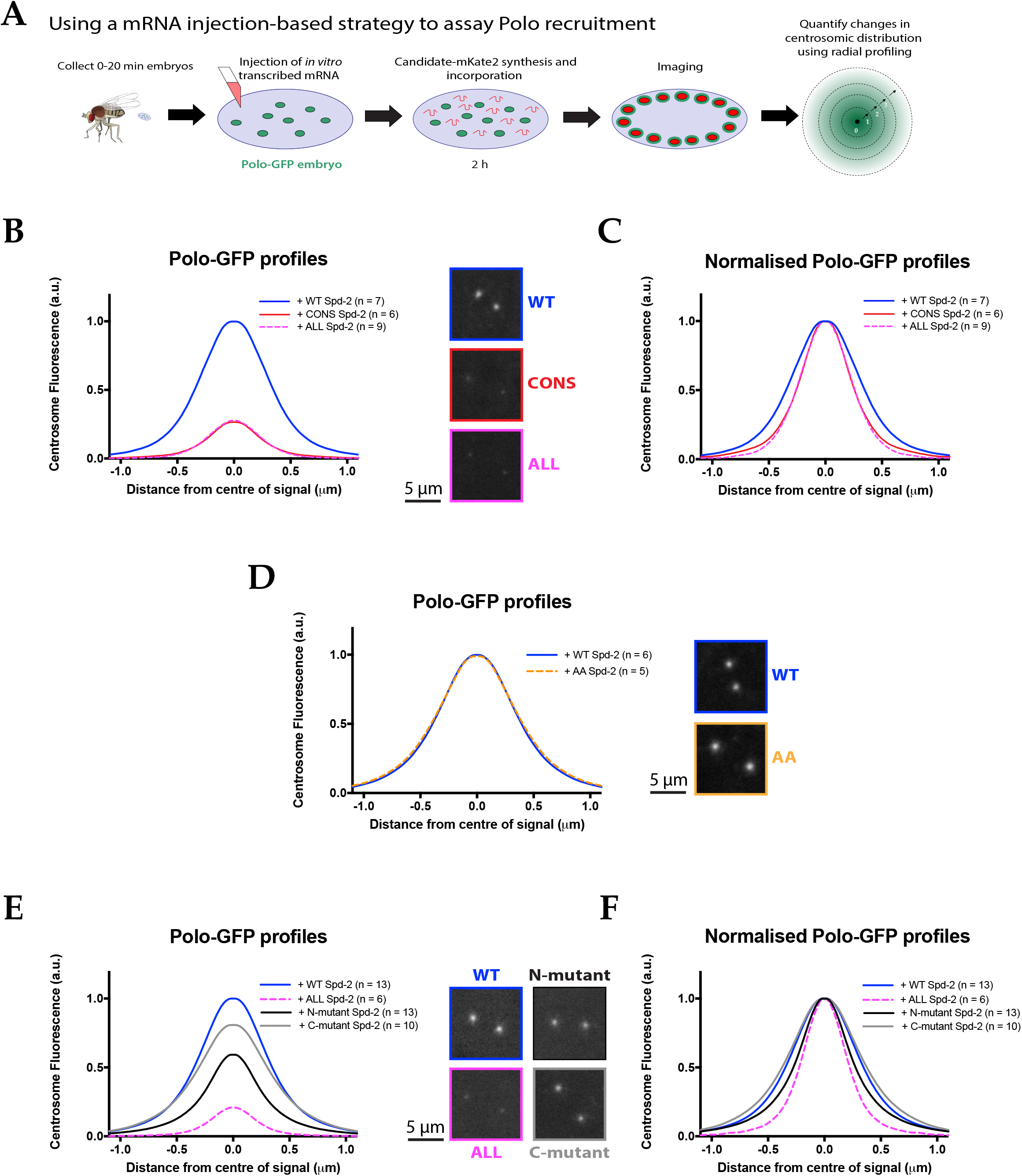
No single site in Spd-2 is required to recruit Polo to centrosomes. **(A)** Schematic illustration of the mRNA injection assay used to analyse the effect of various Spd-2-mKate2 fusion proteins on Polo-GFP recruitment. **(B)** Graph compares the radial distribution of Polo-GFP around the mother centriole in living WT embryos expressing Polo-GFP and injected with mRNAs encoding either WT Spd-2-mKate2, Spd-2-CONS-mKate2 or Spd-2-ALL-mKate2, as indicated. Insets show examples of typical spinning disk confocal images used for this analysis; 6-9 embryos (as indicated), and 5 centrosomes in each embryo, were analysed. **(C)** Graph shows the same data as shown in (B), but normalised so that the peak intensity of all genotypes = 1. This emphasises how even if the centrosomal Polo-GFP signal is normalised for fluorescence intensity, Polo-GFP spreads out around the mother centriole to a lesser extent in the embryos expressing Spd-2-CONS-mKate2 or Spd-2-ALL-mKate2 than WT Spd-2-mKate2. These observations recapitulate our findings from transgenic lines expressing Spd-2-GFP-fusions in a *Spd-2* mutant background. **(D)** Graph compares the radial distribution of Polo-GFP around the mother centriole in living WT embryos expressing Polo-GFP and injected with mRNAs encoding either WT Spd-2-mKate2 or Spd-2-AA-mKate2, as indicated. Insets show examples of typical spinning disk-confocal images used for this analysis; 5-6 embryos (as indicated), and 5 centrosomes in each embryo, were analysed. Note that these datasets are not normalised to each other, and the AA mutation does not detectably perturb the localisation of Polo-GFP. **(E,F)** Same analysis as presented in panels (B,C), but analysing the distribution of Polo-GFP in embryos expressing either Spd-2-ALL-mKate2, or one of two versions of Spd-2 in which the potential Polo binding sites in either the N-terminal or C-terminal region of Spd-2 have been mutated (as indicated).

We first confirmed that we could use this assay to recapitulate our finding that the centrosomal recruitment of Polo-GFP was severely compromised by the expression of Spd-2-CONS-mKate2 or Spd-2-ALL-mKate2. In these embryos, the mutant proteins are gradually translated from the injected mRNA and so they eventually out-compete the endogenous (unlabelled) WT Spd-2 protein (Figure 9B,C). The single S-S(p) motif in SPD-2 that recruits PLK-1 to centrosomes in *C. elegans* is a potential CDK1 substrate that may be conserved in *Drosophila* species *(blue* box, Figure S5). *Drosophila* Spd-2 contains only one other conserved S-S/T motifs that is a potential CDK1 substrate *(yellow* box, Figure S5), so we mutated both of these sites to the non-phosphorylatable residue Ala (T516A and S625A). This construct (Spd-2-AA-mKate2), however, did not detectably perturb the centrosomal distribution of Polo-GFP (Figure 9D).

We then tested whether the ability of fly Spd-2 to recruit Polo to the PCM depended on S-S/T motifs in the N-terminal half of the protein, as all the motifs identified in worms and vertebrates are restricted to this half of the SPD-2/Cep192 molecule (Figure S5). We divided the 34 S-S/T motifs of Spd-2 into 2 equal groups based on their position along the protein (N-terminal or C-terminal) (Figure 2A), and generated constructs with either group of 17 motifs mutated to T-S/T (Spd-2-N-term-mKate2 and Spd-2-C-term-mKate2, respectively). Both constructs led to a reduction in Polo-GFP levels at the centrosome (Figure 9E,F). The mutation of sites in the N-terminal half of Spd-2 had the biggest effect, but this was still mild compared to the injection of Spd-2-ALL-mKate2 mRNA (Figure 9E,F). We conclude that there is no single specific S-S/T(p) motif in Spd-2 that is essential to recruit Polo to the mitotic PCM, and that motifs in both the N- and C-terminal regions contribute to this process.

## Discussion

Centrosome maturation appears to be a near-universal feature of the metazoan cell cycle. Although many of the key proteins required for centrosome maturation have been identified, how these proteins drive this process is unclear. Our data suggests that three proteins—Spd-2, Polo and Cnn—cooperate to form a positive feedback loop that promotes the expansion of the mitotic centrosome in flies (Figure 10A,B).

**Figure 10.**
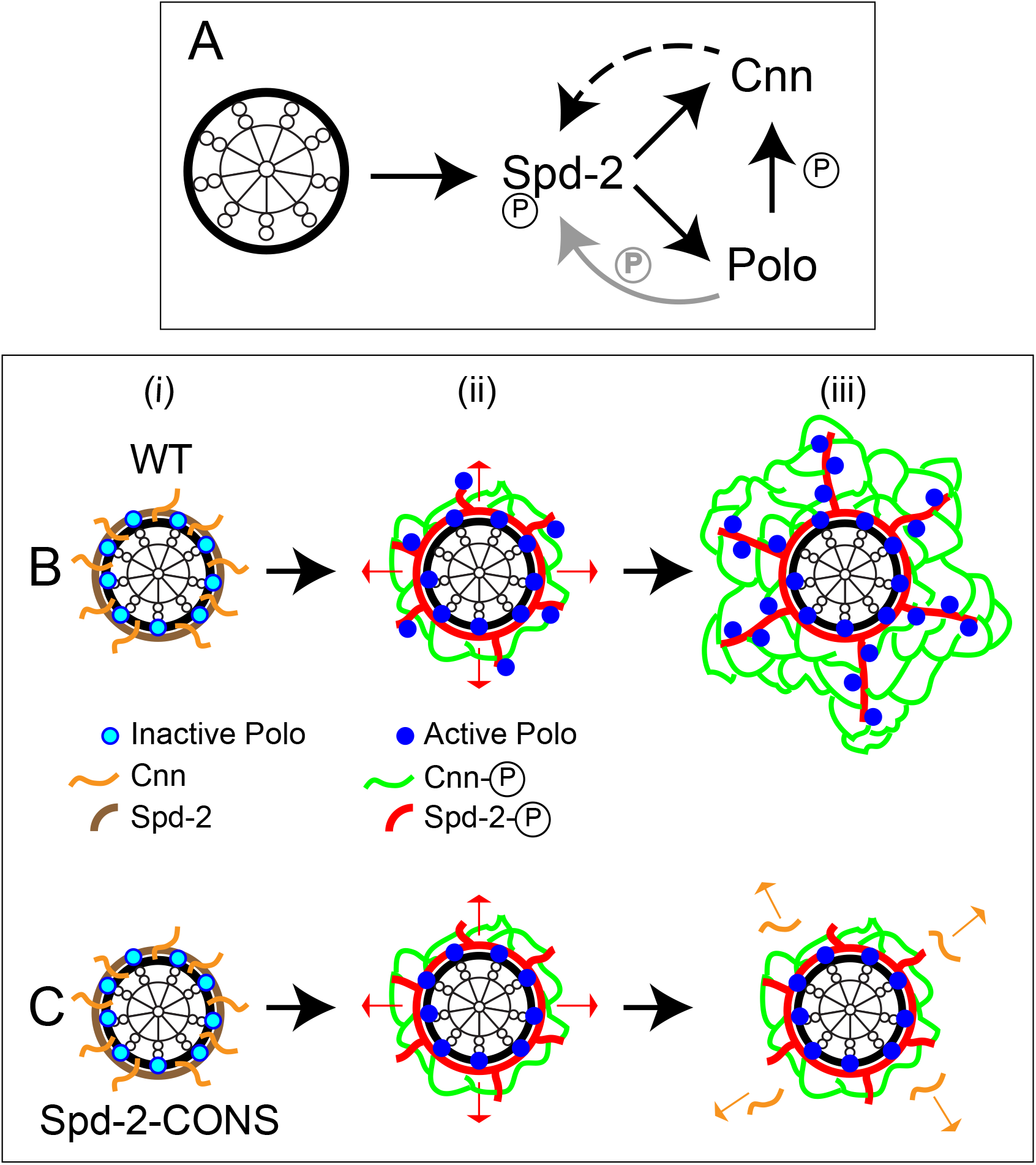
An illustration of the putative Spd-2, Polo and Cnn positive feedback loop. **(A)** A schematic summary of the putative positive feedback loop that drives centrosome maturation in flies. Solid lines indicate recruitment or phosphorylation; dashed line indicates that Cnn does not *recruit* Spd-2, but rather helps to *maintain* Spd-2 that has been recruited by the mother centriole and is fluxing outwards. The grey arrow indicates that Polo may well phosphorylate Spd-2 to create additional S-S/T(p) motifs that can then recruit more Polo, but this is not proven. Note how the Spd-2/Polo/Cnn feedback loop is autocatalytic, but ultimately still relies on the mother centriole as a source of Spd-2, because Cnn itself cannot recruit more Spd-2 or Polo. **(B,C)** Cartoons illustrate the process of centrosome maturation in a WT cell (B), and a cell in which Spd-2 cannot recruit Polo (C). During interphase **(i)**, Spd-2, Polo and Cnn are all recruited to a toroid that surrounds the mother centriole. Polo is inactive, and Spd-2 and Cnn are not phosphorylated. As cells prepare to enter mitosis **(ii)**, Polo is activated, and the centrosomal Spd-2 and Cnn are phosphorylated, allowing them to initially assemble into a “mini-scaffold” around the mother centriole. The phosphorylated Spd-2 scaffold then starts to flux away from the mother centriole (*red arrows*). In normal cells **(B[iii])**, the expanding Spd-2 scaffold recruits more Cnn and more Polo, allowing more Cnn scaffold to assemble. The Cnn scaffold cannot recruit more Spd-2 or Polo, but it stabilises the expanding Spd-2 scaffold, so allowing the Spd-2 scaffold to accumulate around the mother centriole. This creates a positive feedback loop that drives the rapid expansion of the Spd-2 and Cnn scaffolds around the mother centriole, and so centrosome maturation. In fly embryos, the Cnn scaffold also fluxes outwards along the centrosomal MTs (Conduit and Raff, 2015), but this is not illustrated here. If the expanding Spd-2 scaffold cannot recruit Polo **(C[iii])**, the Cnn recruited to the expanding Spd-2 scaffold does not get phosphorylated, and so it cannot form a scaffold. The recruited Cnn rapidly dissipates into the cytosol (*orange* arrows), the positive feedback loop is broken, and centrosome maturation fails.

In fly S2 cultured cells, Spd-2, Polo and Cnn are recruited around the surface of the mother centriole during interphase (Fu and Glover, 2012); Polo is presumably inactive and Spd-2 and Cnn are not phosphorylated, so no pericentriolar scaffold is assembled (Figure 10B[i]). As cells prepare to enter mitosis (Figure 10B[ii]), Polo is activated and the centrosomal Spd-2 becomes phosphorylated (potentially by Polo, but also by other mitotic kinases as well), allowing it to form a scaffold that can recruit both Polo (via phosphorylated S-S/T(p) motifs, as we show here) and Cnn (via an as yet uncharacterised interaction) (Conduit et al., 2014b); this allows Polo to phosphorylate Cnn, which allows Cnn to assemble into its own scaffold structure (Conduit et al., 2014a; Feng et al., 2017). These interactions are sufficient to establish a mini-scaffold around the mother centriole that can recruit some PCM and organise some MTs. However, the phosphorylated Spd-2 scaffold can flux outwards away from the mother centriole (indicated by the *red* arrows, Figure 10B[ii]); this expanding Spd-2 scaffold can recruit more Cnn and, as its phosphorylated, it can also recruit more Polo. The recruited Polo very likely ensures that the Spd-2 scaffold remains phosphorylated, and it also phosphorylates Cnn, allowing the expanding Spd-2 scaffold to continue assembling a Cnn scaffold around itself (Figure 10B[iii]). Crucially, the Cnn scaffold cannot itself recruit Spd-2 or Polo, but it can help to maintain the Spd-2 scaffold that was initially recruited by the mother centriole and is now fluxing outwards. In this way, the Cnn scaffold allows Spd-2—and so Polo—to accumulate around the centriole, which in turn drives more Cnn incorporation into the PCM. This positive feedback mechanism (Figure 10A) could provide the “autocatalysis” that previous mathematical modelling indicates is required to explain the rapid expansion of the mitotic PCM (Zwicker et al., 2014).

If Spd-2 cannot recruit Polo (Figure 10C), it can still recruit Cnn, and this Cnn can still initially be phosphorylated—presumably because it is physically close enough to the pool of Polo that is still present around the mother centriole. Thus, Spd-2, Polo and Cnn can still form the “mini-scaffold” around the mother centriole that can recruit some PCM and organise some MTs (Figure 10C[ii]). The Spd-2 scaffold that assembles, however, cannot recruit any more Polo; as a result, the Cnn recruited by the Spd-2 network is not phosphorylated, and so the mitotic PCM fails to expand around the mother centriole (Figure 10C[iii]).

Importantly, although this proposed mechanism is autocatalytic (because as the Spd-2 scaffold increases in size it can recruit Cnn into the centrosome at a faster rate), it ultimately relies on the mother centriole as a source of Spd-2 (Figure 10A). This neatly solves the conundrum of how mitotic PCM growth appears to be autocatalytic (Zwicker et al., 2014), yet mitotic PCM size is ultimately set by the size of the centriole (Kirkham et al., 2003). For this mechanism to work it is crucial that Cnn is not itself able to efficiently recruit Spd-2 or Polo into the scaffold. If it could do so, mitotic PCM growth would no longer be constrained by the centriole as, once recruited, Cnn could effectively catalyse its own recruitment (by recruiting more Spd-2 and Polo). Spd-2 and Cnn are of similar size in flies (1146aa and 1148aa, respectively) but Spd-2 has >5X more conserved potential PBD-binding (S-S/T) motifs than Cnn (Figure S6). Interestingly, a similar ratio is found when comparing human Cep192 (1941aa) to human Cep215/Cdk5Rap2 (1893aa) (Figure S6), even though the human and fly homologues of both proteins share only very limited amino acid identity. We speculate that these two protein families may have evolved to ensure that phosphorylated Spd-2/Cep192 can efficiently recruit Polo/Plk1, whereas phosphorylated Cnn/Cep215 cannot.

Another important aspect of this model is that Spd-2 is incorporated into the mitotic PCM only at the centriole surface, and then fluxes outwards (Conduit et al., 2014b). This Spd-2-flux has so far only been observed in *Drosophila* (Conduit and Raff, 2015; Conduit et al., 2010; 2014b). In fly embryos, Cnn also fluxes outwards but, unlike Spd-2, this flux requires MTs and is only observed in embryos (Conduit and Raff, 2015). In *C. elegans* embryos, SPD-5 behaves like Cnn in somatic cells: it does not flux outwards and is incorporated isotropically throughout the volume of the PCM (Laos et al., 2015). Clearly it will be important to determine whether Spd-2/Cep192 homologues exhibit a centrosomal-flux in worms and other non-fly species, and whether this flux provides the primary mechanism by which the mother centriole continuously influences the assembly of the expanding mitotic PCM.

In vertebrates, Cep192 serves as a scaffold for Plk1 and also Aurora A (Joukov et al., 2010; 2014; Meng et al., 2015)—another mitotic protein kinase that plays an important part in centrosome maturation in many species (Barr and Gergely, 2007). Here, there appears to be a complex interplay between Cep192, Plk-1 and Aurora A, with Cep192 acting as a scaffold that allows these two important regulators of mitosis to influence each other’s activity and centrosomal localisation. Spd-2 clearly plays an important part in recruiting Aurora A to centrosomes in fly cells (Conduit et al., 2014b; Dobbelaere et al., 2008)—although it is unclear if this is direct, as fly and worm Spd-2/SPD-2 both lack the N-terminal region in vertebrate Cep192 that recruits Aurora A (Meng et al., 2015) (Figure S5). How Aurora A might influence the assembly of the Spd-2, Polo/PLK-1 and Cnn/SPD-5 scaffold in flies and worms remains to be determined.

Finally, there has been great interest recently in the idea that many non-membrane bound organelles like the centrosome may form by phase separation and have liquid like properties (Banani et al., 2017; Rousseau et al., 2018). In support of this possibility for the centrosome, purified recombinant SPD-5 can assemble into condensates *in vitro* that have transient liquid-like properties, although they rapidly harden into a more viscous gel- or solid-like phase (Woodruff et al., 2016). Moreover, a mathematical model that accurately describes centrosome maturation in the early worm embryo treats the centrosome as a liquid, and it is from this model that the importance of autocatalysis was first recognised (Zwicker et al., 2014). *In vivo*, however, the Cnn and SPD-5 scaffolds do not appear to be very liquid-like (Conduit et al., 2010; Conduit et al., 2014a; 2014b; Laos et al., 2015) and fragments of Cnn can assemble into micron-scale assemblies *in vitro* that are clearly solid- or viscous-gel-like (Feng et al., 2017). Our data suggests that the incorporation of Spd-2 into the PCM only at the surface of the centriole, coupled to an amplifying Spd-2/Polo/Cnn positive feedback loop, could provide an “autocatalytic” mechanism that functions within the conceptual framework of a non-liquid-like scaffold that emanates from the mother centriole.

